# Mental Health, Schooling Attainment and Polygenic Scores: Are There Significant Gene-Environment Associations?

**DOI:** 10.1101/684688

**Authors:** Vikesh Amin, Jere R. Behrman, Jason M. Fletcher, Carlos A. Flores, Alfonso Flores-Lagunes, Hans-Peter Kohler

## Abstract

It is well-established that (1) there is a large genetic component to mental health, and (2) higher schooling attainment is associated with better mental health. Given these two observations, we test the hypothesis that schooling may attenuate the genetic predisposition to poor mental health. Specifically, we estimate associations between a polygenic score (PGS) for depressive symptoms, schooling attainment and gene-environment (GxE) interactions with mental health (depressive symptoms and depression), in two distinct United States datasets at different adult ages-29 years old in the National Longitudinal Study of Adolescent Health (Add Health) and 54 years old in the Wisconsin Longitudinal Study (WLS). OLS results indicate that the association of the PGS with mental health is similar in Add Health and the WLS, but the association of schooling attainment is much larger in Add Health than in the WLS. There is some suggestive evidence that the association of the PGS with mental health is lower for more-schooled older individuals in the WLS, but there is no evidence of any significant GxE associations in Add Health. Quantile regression estimates also show that in the WLS the GxE associations are statistically significant only in the upper parts of the conditional depressive symptoms score distribution. We assess the robustness of the OLS results to omitted variable bias by using the siblings samples in both datasets to estimate sibling fixed-effect regressions. The sibling fixed-effect results must be qualified, in part due to low statistical power. However, the sibling fixed-effect estimates show that college education is associated with fewer depressive symptoms in both datasets.

## 1. Introduction

Mental illness is an important public health issue in the US because of its high prevalence and economic and societal costs. In any given year, 18.1% of adults aged 18 or older suffer from some mental illness (Center for Behavioral Health Statistics and Quality 2015). Serious mental illnesses cost $193.2 billion in lost earnings (Insel 2008) and reduce life expectancy by 10-20 years (Chesney et al. 2014). Suicide is the 10^th^ leading cause of death in the US and one of the factors driving the recent fall in life expectancy (Murphy et al. 2018).

Mental illness is a complex health condition. There is increasing evidence showing that interactions between environmental and genetic factors (gene-environment interactions; GxE) contribute to poor mental health. The first study to show this was Caspi et al. (2002). They found that associations between childhood maltreatment and antisocial behavior were modified by the MAOA gene, with those holding the low-activity variant being more responsive to maltreatment than the high-activity group. In a second paper, Caspi et al. (2003) found that a functional polymorphism in the 5-HTT gene moderated the influence of stressful life events on depression. Since the publication of these findings, several studies have looked at interactions between childhood maltreatment/stressful life events and candidate genes (MAOA, COMT, DRD4, BDNF, 5-HTTLPR) that predict poor mental health.

However, candidate genes are rarely used now to measure genetic predisposition. Candidate genes only explain a small proportion of the variation in mental health (Duncan and Keller 2011), whereas the genetic architecture of mental health is highly polygenic (involves multiple genes). Candidate gene studies are therefore underpowered, and previous significant candidate GxE associations are likely to be false positives (Border et al. 2019).^1^ The current best practice in the literature is to use a polygenic score (PGS), which is a summary measure of an individual’s genetic predisposition for a given trait. More specifically, a PGS is a weighted count of the number of trait-associated alleles possessed by an individual. The weights are based on effect sizes from genome-wide association studies (GWAS), conducted in large independent samples. PGSs can be limited to genome-wide significant single nucleotide polymorphisms (SNPs) (p< 5×10^-8^), or a series of more liberal thresholds can be applied (e.g., p<.0.001, 0.01). Studies using PGSs (for major depressive disorder; MDD) have reached mixed conclusions regarding GxE interactions with respect to childhood trauma and stressful life events. Peyrot et al. (2014) found that individuals with a high PGS, and a history of childhood trauma were more likely to develop MDD than those with a low PGS and no exposure to trauma. Mullins et al. (2016) also found a significant GxE association between the PGS and childhood trauma. However, the GxE association was in the opposite direction-individuals who experienced moderate-to-severe childhood trauma had a lower PGS.^2^ Both Mullins et al. (2016) and Musliner et al. (2016) found insignificant associations between the PGS and stressful life events in adulthood.

In this paper, we examine interactions between genes and another environmental factor, schooling attainment, which to our knowledge has not previously been examined in the GxE interaction literature for mental health. Several empirical studies have shown that individuals with more schooling have better mental health and are less likely to be depressed (see Lorant et al. 2003 for a meta-analysis).^3^ There are several theoretical reasons for this relationship. First, schooling can directly improve health by increasing productive efficiency (producing more health from a given set of input) and allocative efficiency (choosing better inputs) (Grossman 1972). Schooling also improves mental health because it is associated with fewer and less severe stressors. For example, higher schooling attainment is associated with higher wages (Card 1999), which reduce financial stress and provide the means to pursue better mental health. Moreover, higher schooling attainment is associated with lower risks of unemployment and work-related adverse life events, which can lead to poor mental health (Audhoe et al. 2010). More schooling is further associated with leading healthier lifestyles (Cutler and Lleras-Muney 2010), and interacting with other healthy peers (Gaviria and Raphael 2001), which all positively affect mental health. More generally, schooling is called a fundamental cause of health disparities because of the perception that schooling embodies access to important resources and affects multiple health outcomes through several mechanisms (Link and Phlean 1995).

Given the apparent importance of schooling, we hypothesize that schooling may attenuate the genetic predisposition to poor mental health. There are theoretical models suggesting that schooling may interact with genes. The diathesis-stress model (Ellis et al. 2011) hypothesizes that unhealthy environments (low schooling attainment) trigger risk alleles, while healthy environments (high schooling attainment) protect against risk alleles. For example, a college graduate with a high genetic risk of depression may never be depressed because being a college graduate is associated with higher income, leading a healthier lifestyle, and interacting more with other healthy peers. In contrast, a high school dropout with a high genetic risk is more likely to be depressed because he/she experiences financial stress due to having a low income or being unemployed, leads an unhealthy lifestyle and interacts with other depressed peers. The differential susceptibility (also commonly known as the orchids and dandelions) model (Belsky and Pluess 2009) hypothesizes that some individuals are dandelions, meaning they have a genetic make-up that is unaffected by both positive and negative environments. Others are orchids whose genetic make-up is highly sensitive to the environment. These individuals thrive in positive environments but wilt in negative environments. This model therefore predicts that schooling is more likely to lower the influence of genes for orchid individuals who have a high genetic predisposition of depression.

We examine the hypothesis that gene-education interactions are important for mental health by using OLS regressions to estimate associations between genes, schooling attainment and GxE interactions with mental health for European-ancestry individuals in the National Longitudinal Study of Adolescent Health (Add Health) and the Wisconsin Longitudinal Study (WLS).^4^ We analyze mental health when individuals are 29 years old in Add Health and 54 years old in the WLS. We measure mental health using the Center for Epidemiologic Studies Depression (CES-D) Scale in both datasets. The CES-D is widely used to measure symptoms of depression, and has been tested in multiple settings for validity and reliability (Cosco et al. 2017; Radloff 1977). We use a PGS for the number of depressive symptoms created by the Social Science Genetic Association Consortium (SSGAC) in both datasets. Both datasets also oversample siblings, which allows us to estimate sibling fixed-effect regression models. Sibling fixed-effect models control for (i) observed and unobserved shared family-level factors including shared childhood neighborhoods and schools and (ii) unobserved shared portions of the genome that affect schooling and mental health – both of which may bias OLS estimates. On the other hand, sibling fixed-effects do not eliminate sibling-specific differences that affect schooling and mental health, and thus may still yield biased estimates of causal effects. However, sibling fixed-effects estimates are probably still an improvement on OLS estimates that do not control for any confounding (other than the regressors included in the model).

To our knowledge, this is the first study to estimate associations between a PGS for depressive symptoms, schooling attainment and GxE interactions with mental health. The closest studies related to our paper are Amin et al. (2017), Liu et al. (2015), and Komulainen et al. (2018). These studies provide OLS associations between a PGS for BMI, schooling attainment and GxE interactions with BMI and obesity. The general conclusion from these studies is that there are significant associations of the PGS and schooling attainment, but no strong evidence of significant GxE associations.^5^

The comparison between Add Health and the WLS is interesting for at least two reasons. First, schooling-health gradients may differ over the life cycle. The cumulative advantage hypothesis (Ross and Wu 1996) predicts that schooling-health gaps emerge by early adulthood and widen over the life cycle as the benefits of schooling accumulate. The age-as-leveler hypothesis (Herd 2006) predicts that schooling-health gaps narrow over the life cycle, as the importance of biological determinants increase relative to the importance of schooling. The empirical evidence is consistent with the cumulative advantage hypothesis until middle age, and with biological determinants being more important in old age.^6^ If schooling-mental health gaps are smaller at older ages, then one might expect smaller GxE interactions in the WLS than in Add Health. Second, schooling is left-truncated at 12 grades in the WLS as it follows a sample of high school graduates. Therefore, there could be smaller schooling-mental health gradients in the WLS, and smaller GxE interactions.

Our OLS results indicate that the association of the PGS with mental health is similar in Add Health and the WLS, but the association of schooling attainment is much larger in Add Health than in the WLS. We find no evidence of any significant GxE associations in Add Health, but there is some suggestive evidence that the association of the PGS with mental health is lower for more-schooled older individuals in the WLS. Quantile regression estimates also show that in the WLS the GxE associations are statistically significant only in the upper parts of the conditional depressive symptoms score distribution, that is, for individuals who exhibit higher levels of depression than the average person in the data. It is difficult to draw firm conclusions from the sibling fixed-effect results, in part due to low statistical power. However, the sibling fixed-effects estimates do show that college education is associated with fewer depressive symptoms in both datasets.

The paper is organized as follows. We describe the datasets and the methodology in sections 2 and 3, respectively. We present and discuss the results in section 4. Finally, section 5 concludes.

## 2. Data

### 2.1 The National Longitudinal Study of Adolescent Health (Add Health)

Add Health is a nationally-representative sample of 20,745 students in grades 7 through 12 (aged 12-21) in 1994-95 (wave 1). Adolescents were surveyed from 132 schools that were selected to ensure representativeness with respect to region, urbanicity, school size and type, and ethnicity. In wave 1, data were collected from adolescents, their parents, siblings, friends, relationship partners, fellow students, and school administrators. The adolescents have been followed after 1 year (wave 2, 1996), 6 years (wave 3, 2001-2002), 13 years (wave 4, 2008), and 21 years (wave 5, 2016-2018). Add Health includes oversamples of ethnic minorities, disabled students, and sibling pairs living in the same household.

We examine the relationship between genetics, schooling attainment, and mental health at wave 4.^7^ Mental health is measured using the 10 item CES-D score. The CES-D score is created by summing responses (ranging from 0 to 3) to questions that asked respondents how often in the last week they (1) were bothered by things not normally bothersome; (2) could not shake the blues; (3) felt like they were not as good as others; (4) had trouble focusing; (5) were depressed; (6) were too tired to do things they enjoyed; (7) felt sad; (8) felt happy; (9) enjoyed life, and (10) felt disliked. The CES-D score ranges from 0 to 30, with higher values corresponding to poorer mental health. Depression is defined as having a score of 11 or higher (Suglia et al. 2015). Schooling attainment is based on responses to the question “what is the highest level of education that you have achieved to date?” Response options and their assigned grades of schooling (in parentheses) were: eighth grade or less (8), some high school (10), high school graduate (12), some vocational/technical training (13), completed vocational/technical training (14), some college (14), completed college (16), some graduate school (17), completed a master’s degree (18), some graduate training beyond a master’s degree (19), completed a doctoral degree (20), some post-baccalaureate professional education (18), and completed post-baccalaureate professional education (19).

At wave 4 96% of participants consented to providing saliva samples. Approximately 12,200 (80% of those participants) consented to long-term archiving and were consequently eligible for genome-wide genotyping. Genotyping was done on two Illumina platforms, with approximately 80% of the sample genotyping performed with the Illumina Omni1-Quad BeadChip and 20% genotyped with the Illumina Omni2.5-Quad BeadChip. After quality-control procedures, genotyped data are available for 9,974 individuals (7,917 from the Omni1 chip and 2,057 from the Omni2 chip) on 609,130 SNPs common across both genotyping platforms.

The SSGAC has created a PGS for depressive symptoms using the genetic data. Briefly, the PGS was constructed using LDpred, a Bayesian method that includes all measured SNPs and weights each SNP by an approximation to its conditional effect, given other SNPs.^8^ The PGS was calculated with data from all SNPs; no statistical significance threshold was applied to restrict the SNPs that were included. The weights for the PGS are based on Multi-Trait Analysis of GWAS (MTAG; Turley et al. 2018) summary statistics. MTAG is a method that uses GWAS summary statistics for a primary phenotype (depressive symptoms in our case) and for one or more secondary phenotypes (neuroticism and subjective well-being in our case) to produce an updated set of summary statistics for the primary phenotype that, under certain assumptions, will be more precisely estimated than the input GWAS summary statistics.^9^ A PGS constructed using MTAG summary statistics has more predictive power than a PGS that employs standard GWAS summary statistics (Turley et al. 2018).

The SSGAC provides the standardized PGS for 5,690 European ancestry respondents.^10^ Our final sample consists of 5335 European ancestry individuals with non-missing information on the PGS, mental health, schooling attainment and the control variables. The control variables are gender, age, birth order, mother’s schooling attainment, adolescent IQ, and a PGS for educational attainment constructed by the SSGAC using the MTAG method, which we use to control for unobserved ability. We measure mother’s schooling attainment using the schooling attainment of the resident mother from the wave 1 questionnaire. In order to maximize the sample size, missing mother’s schooling is imputed with the sample mean, and controlled for by using a dummy variable for missing mother’s schooling^11^. Adolescent IQ is measured using the wave 1 Peabody Picture Vocabulary test^12^. Our sibling analyses are based on 457 full biological sibling pairs with non-missing information on the key variables.

### 2.2 The Wisconsin Longitudinal Study (WLS)

The WLS is a longitudinal study of a one-third random sample of the graduating high school class of 1957 in Wisconsin. Survey information from 10,317 of the graduates was collected at age 18 in 1957. Graduates were followed with a combination of in-person, telephone, and mail surveys at ages 36 (1975), 54 (1993-1994), 65 (2003-2004), and 72 years (2011). The WLS sample also includes information from a randomly selected sibling (who may or may not be a high school graduate) that was collected in 1977, 1993-1994, 2005-2007, and 2011. The sample is broadly representative of white, non-Hispanic American men and women who have completed at least a high school education in Wisconsin. In total, 19,050 graduates and their siblings have contributed data.

We examine the relationship between genetics, schooling attainment, and mental health in 1992-1994, when mental health was first measured. Mental health is measured using the 20 item CES-D score. The scoring method in the WLS differs from the standard CES-D scoring procedure. Respondents were asked for the number of days they experienced a particular event, with options ranging from 0 to 7 days. This was converted to the standard CES-D scoring procedure by coding 0 days as 0, 1-2 days as 1, 3-4 days as 2, and 5-7 days as 3. The CES-D scale thus ranges from 0-60. Depression is defined as having a CES-D score of 22 or higher for men and 24 or higher for women (Roberts et al. 1991). Schooling attainment is measured from the 1975-1977 surveys. The WLS assigns 12 grades for graduating high school; 13 for one year of college; 14 for an associates degree; 15 for college trained nurses; 16 for a bachelors degree; 17 for a masters degree; 18 for a two-year masters degree; 19 for one or more years post the two-year masters degree, and 20 for a doctorate degree.

The WLS first collected saliva samples in 2007-2008 by mail. Additional samples were collected in the course of home interviews in 2010. A total of 9472 study samples were successfully genotyped at the Center for Inherited Disease Research (CIDR) at Johns Hopkins University. Genotyping was performed at CIDR using the Illumina HumanOmniExpress array and using the calling algorithm GenomeStudio version 2011.1, Genotyping Module version 1.9.4, GenTrain Version 1.0. After quality control, genetic data were available for 9,012 respondents. Of these respondents, 64% are graduate members of the sample and 36% are siblings of graduates. The SSGAC has created a PGS for depressive symptoms using the same procedure that was used with the Add Health genetic data.

The PGS is available for 8,509 respondents (5,414 graduates and 3,095 siblings). Our final sample consists of 5,521 European ancestry individuals with non-missing information on the PGS, mental health, schooling attainment and the control variables.^13^ The control variables are gender, age, birth order, mother’s schooling attainment, adolescent IQ, and a PGS for educational attainment constructed by the SSGAC using the MTAG method. Mother’s schooling attainment comes from the graduate respondents’ responses to the 1957 questionnaire. Mother’s schooling attainment is missing for siblings, who were not asked about their mother’s schooling. For siblings we impute mother’s schooling with the mother’s schooling reported by the graduate, and control for an indicator for missing mother’s schooling. Adolescents IQ is based on the Henmon-Nelson test of mental ability, which is a group-administered, 30-min, multiple-choice assessment that consists of 90 verbal or quantitative items and that all high school students in Wisconsin were required to take in their third year of high school. Our sibling analyses are based on 1114 full biological sibling pairs with non-missing information on the key variables. We expect to have more statistical power with these data relative to the sample of siblings from Add Health given the larger sample size.

## 3. Methodology

We estimate associations between genetics, schooling, and GxE interactions with mental health through OLS regressions based on equation (1) in which the mental health outcome of individual *i* (*MH*_*i*_) is related to a PGS for depressive symptoms (*PGS*_*i*_), grades of schooling (*Schooling*_*i*_), an interaction between the PGS and grades of schooling (*PGS*_*i*_ * *Schooling*_*i*_), a set of control variables noted above (*X*_*i*_), and an error term (*u*_*i*_).

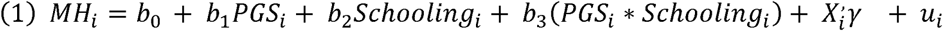

Although genes are randomly inherited from parents at conception, *b*_1_ and *b*_3_ could be biased because of population stratification. Population stratification refers to a situation where the distribution of genes systematically differs by population subgroups (e.g. by ethnicity/race). If these subpopulations also systematically have different health outcomes that are not due to genetic make-up, then this could lead to a spurious correlation between the PGS and health. Population stratification can be controlled by limiting analyses to ethnically homogenous samples (Cardon and Palmer 2003) and by including principal components from genome-wide SNP data as control variables, which account for genetic differences across ethnic groups (Price et al. 2006). We include the first 10 principal components to control for population stratification and limit our analyses to individuals of European-ancestry. Therefore, our analysis assumes the variation in PGS is exogenous conditional on population stratification.

Even though we add as regressors variables related to the individual’s ability (e.g. adolescent IQ and PGS for educational attainment) and family background (e.g., maternal education), schooling attainment may still be correlated with unobservable confounders, in which case *b*_2_ and *b*_3_ would be biased. To reduce possible unobserved confounding, we estimate sibling fixed-effects regressions (equation 2) where the mental health outcome of sibling i in pair j (*MH*_*ij*_) is related to the PGS (*PGS*_*ij*_), grades of schooling (*Schooling*_*ij*_), an interaction between the PGS and grades of schooling (*PGS*_*i*_ * *Schooling*_*ij*_), individual controls (*X*_*ij*_), sibling fixed effects (dummy variables for each sibling pair, μ_*j*_), and an error term (*ε*_*ij*_).

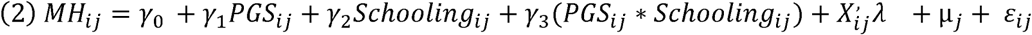

Siblings share about 50% of their genes if mating is random (more if mating is positively assortative) and the family environment is very similar. This means that the inclusion of sibling fixed-effects controls for (i) population stratification, (ii) observed and unobserved family-level factors, and (iii) about 50% (or more) of unobserved genetic factors that affect schooling and mental health. The variation is considered to be quasi-exogenous because differences in genotypes of full biological siblings are the outcomes of a genetic lottery (Fletcher and Lehrer 2011). Sibling fixed-effects estimates do not eliminate unobserved sibling-specific differences, and so may still be biased. Nevertheless, comparisons between OLS and sibling fixed-effects may provide information about the extent to which OLS estimates are biased because of a failure to account for confounding due to the shared genes and family environment for which siblings fixed-effects estimates control.

## 4. Results

### 4.1 Descriptive Statistics

Summary statistics for the Add Health and WLS estimation samples are in columns 1 and 4 in table 1. Add Health respondents are 29 years old on average, and WLS respondents are 54 years old. The proportion of women in both datasets is similar (53% in Add Health; 55% in the WLS). Add Health respondents have higher schooling attainment on average. They have a mean of 14.6 grades of schooling, compared to 13.8 grades for WLS respondents. The main difference is that a higher proportion of individuals have some post high-school education in Add Health. In Add Health, 44% have some (but not complete) college education and 33% are college graduates. In comparison, in the WLS only 15% have some (but not complete) college education and 30% are college graduates. Average mother’s schooling is also higher in Add Health (13.4 grades) than in the WLS (10.7 grades). The average CES-D score in Add Health is 5.88 and 10.55 in the WLS. Recall, that the CES-D scores are not directly comparable, because in Add Health the CES-D score is based on 10 items and ranges from 0-30, while the CES-D score in the WLS is based on 20 items and ranges from 0-60. A higher proportion of individuals are classified as being depressed in Add Health (15%) compared to the WLS (8%). Individuals that consented to provide genetic data may be different from those that did not give consent. In columns 2 and 5 we provide summary statistics for white individuals in Add Health and the WLS who have non-missing information on the key variables and do not have any genetic data. The summary statistics show that white individuals who do not have any genetic data are very similar in terms of observed characteristics to those who are in our estimation samples.

**Table 1:** Summary Statistics.

|  | Add Health<br>Estimation<br>Sample | Add Health<br>Whites<br>Sample with<br>no genetic data | Add Health<br>Sibling<br>Sample | WLS<br>Estimation<br>Sample with<br>no genetic data | WLS<br>Whites<br>Sample | WLS<br>Sibling<br>Sample |
| --- | --- | --- | --- | --- | --- | --- |
|  | (1) | (2) | (3) | (4) | (5) | (6) |
| **Basic Demographics** |  |  |  |  |  |  |
| Age | 28.98 (1.73) | 28.96 (1.72) | 29.00 (1.69) | 53.12 (3.24) | 53.63 (4.37) | 53.09 (4.69) |
| Female | 0.53 (0.50) | 0.53 (0.50) | 0.53 (0.50) | 0.55 (0.50) | 0.55 (0.50) | 0.55 (0.50) |
| **Mental Health** |  |  |  |  |  |  |
| CES-D Score | 5.88 (4.70) | 5.75 (4.59) | 5.97 (4.98) | 10.55 (8.80) | 11.33 (9.47) | 10.49 (8.43) |
| Depressed | 0.15 (0.36) | 0.14 (0.35) | 0.15 (0.36) | 0.08 (0.28) | 0.10 (0.31) | 0.08 (0.27) |
| PGS for depressive symptoms | 0.00 (1.00) | --- | 0.00 (1.00) | 0.00 (1.00) | --- | 0.00 (1.00) |
| **Schooling Attainment** |  |  |  |  |  |  |
| Grades of Schooling | 14.56 (2.18) | 14.89 (2.18) | 14.46 (2.24) | 13.76 (2.33) | 13.44 (2.18) | 13.91 (2.40) |
| High school dropout | 0.07 (0.26) | 0.06 (0.23) | 0.09 (0.28) | 0.01 (0.07) | 0.01 (0.10) | 0.01 (0.10) |
| High school graduate | 0.16 (0.37) | 0.14 (0.35) | 0.17 (0.38) | 0.55 (0.50) | 0.60 (0.49) | 0.51 (0.50) |
| Some college | 0.44 (0.50) | 0.41 (0.49) | 0.42 (0.49) | 0.15 (0.36) | 0.16 (0.36) | 0.16 (0.37) |
| College graduate | 0.33 (0.47) | 0.39 (0.49) | 0.32 (0.47) | 0.30 (0.46) | 0.23 (0.42) | 0.33 (0.47) |
| Education PGS | 0.00 (1.00) | --- | 0.00 (1.00) | 0.00 (1.00) | --- | 0.00 (1.00) |
| **Individual and Family Background Characteristics** |  |  |  |  |  |  |
| IQ | 104.39 (11.73) | 105.50 (12.45) | 103.97 (11.42) | 103.93 (14.68) | 101.56 (15.12) | 104.96 (14.78) |
| Birth order | 1.77 (1.04) | 1.71 (0.99) | 2.02 (1.11) | 2.36 (1.65) | 2.32 (1.63) | 2.43 (1.61) |
| Mother's grades of schooling | 13.39 (1.97) | 13.57 (2.01) | 13.36 (1.85) | 10.73 (2.96) | 10.72 (2.97) | 10.79 (3.02) |
| Mother's schooling missing | 0.09 (0.28) | 0.08 (0.28) | 0.07 (0.20) | 0.23 (0.42) | 0.36 (0.48) | 0.50 (0.50) |
| **Within-Sibling Differences** |  |  |  |  |  |  |
| % of sibling pairs discordant on |  |  |  |  |  |  |
| Depression |  |  | 24% |  |  | 15% |
| College graduate |  |  | 22% |  |  | 30% |
| Absolute mean within-sibling difference |  |  |  |  |  |  |
| In grades of schooling |  |  | 0.82 (0.84) |  |  | 1.85 (2.04) |
| In CES-D Score |  |  | 2.49 (2.18) |  |  | 8.23 (7.70) |
| N | 5335 | 2779 | 914 | 5521 | 3344 | 2228 |

Notes: Standard deviations in parentheses.

Summary statistics for the Add Health and WLS sibling samples are given in columns 3 and 6, respectively. There are two key observations. First, the sibling samples are very similar to the full samples in terms of observed characteristics. For example, in Add Health (WLS, respectively) the average CES-D score for siblings is 5.97 (10.49), 15% (8%) are depressed, and 32% (33%) are college graduates. These summary statistics are similar to those for the estimation sample in column 1 (4). Figures 1 and 2 provide histograms of the CES-D score and depressive symptoms PGS for the sibling and main estimation samples. The distributions of the CES-D score and PGS in the sibling samples are very similar to those in the estimation samples in both datasets. The summary statistics and figures 1 and 2 suggest that the siblings samples may be random samples of the corresponding surveys’ populations. Second, there is a fair amount of within-sibling variation in mental health and schooling attainment that is used to identify the sibling fixed-effects estimates. In Add Health (WLS) 22% (30%) of the sibling pairs are discordant on college graduation, and the absolute mean within-sibling difference in grades of schooling is 0.8 (1.9) grades. In terms of within-sibling differences in mental health 24% (15%) of sibling pairs are discordant on depression in Add Health (WLS), and the absolute mean within-sibling difference in the CES-D score is 2.49 (8.23) units. In terms of sample sizes, the Add Health siblings samples contains 914 siblings (457 sibling pairs) compared to 2228 sibling (1114 sibling pairs) in the WLS. We thus expect to have more statistical power using the WLS siblings sample.

**Figure 1:**
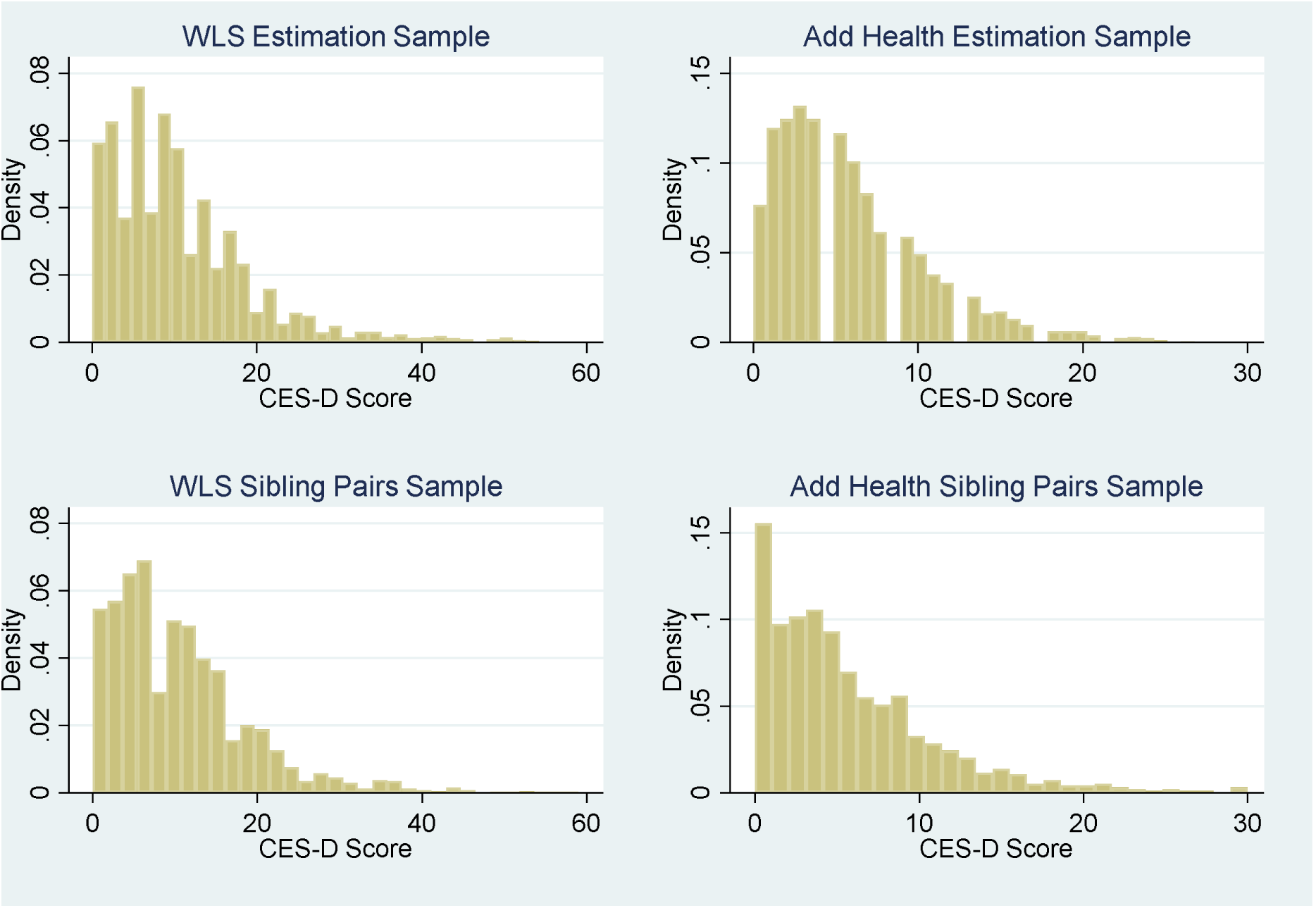
Histogram of CES-D Score.

**Figure 2:**
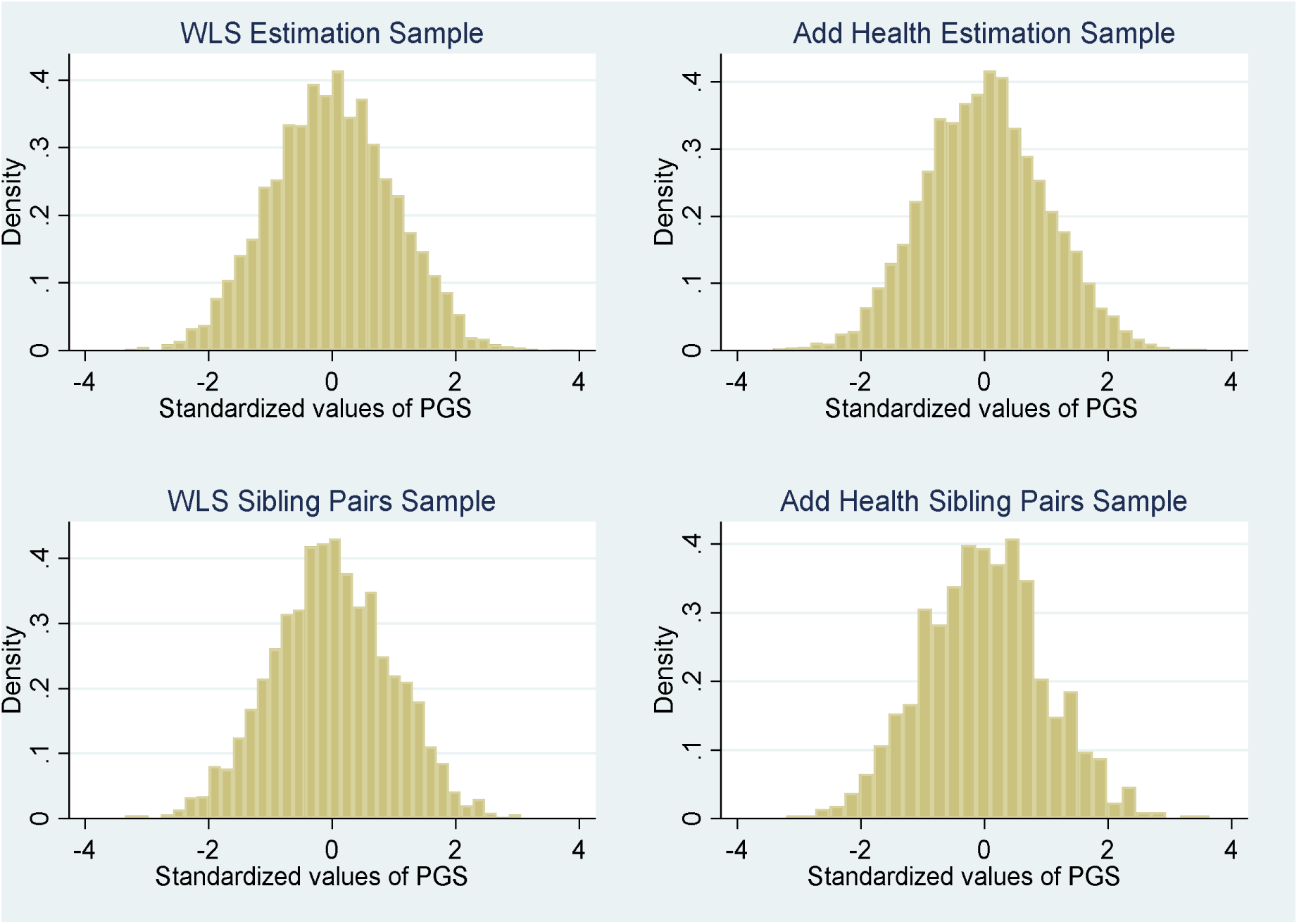
Histogram of PGS for Depressive Symptoms.

### 4.2 OLS Results

The main results for the CES-D score are presented in table 2. Estimates for Add Health are in columns 1-4, and estimates for the WLS are in columns 5-8. In order to compare estimates from both datasets, the CES-D score has been standardized to have a mean of 0 and standard deviation of 1. Estimates in columns 1 and 5 show that the associations of the PGS with depressive symptoms in young adulthood and at older ages are very similar. A 1 standard deviation increase in the PGS is associated with a 0.119 and 0.090 standard deviation increase in the CES-D score in Add Health and the WLS, respectively. Columns 2 and 6 add grades of schooling. More schooling is associated with a much larger decrease (almost 4 times) in depressive symptoms in young adulthood than at older ages. Conditional on the PGS, an additional grade of schooling is associated with a 0.095 (0.025) standard deviation decrease in the CES-D score in Add Health (the WLS). The regression specification in columns 3 and 7 is the same as that in columns 2 and 5, but also controls for birth order, mother’s schooling attainment, adolescent IQ, and the education PGS. Adding these controls decreases the association of grades of schooling slightly in Add Health to 0.085. In contrast, in the WLS the association of grades of schooling drops substantially to −0.006 and is statistically insignificant. Columns 4 and 8 add the interaction between the grades of schooling and the PGS. The coefficient estimates of the interaction terms are statistically insignificant in both datasets.

**Table 2:** OLS Associations of the PGS, Grades of Schooling, and GxE Interactions with the CES-D Score in Add Health and the WLS.

|  | **Add Health** |  |  |  | **WLS** |  |  |  |
| --- | --- | --- | --- | --- | --- | --- | --- | --- |
|  | **(1)** | **(2)** | **(3)** | **(4)** | **(5)** | **(6)** | **(7)** | **(8)** |
| **Genetics** |  |  |  |  |  |  |  |  |
| PGS | 0.119*** | 0.097*** | 0.098*** | 0.083*** | 0.090*** | 0.087*** | 0.093*** |  |
| 0.108*** | (.013) | (.013) | (.013) | (.021) | (.014) | (.014) | (.014) | (.018) |
| **Schooling** |  |  |  |  |  |  |  |  |
| Grades of schooling |  | -0.095*** | -0.085*** | -0.085*** |  | -0.025*** | -0.006 | -0.006 |
|  |  | (.006) | (.007) | (.007) |  | (.006) | (.007) | (.007) |
| **GxE** |  |  |  |  |  |  |  |  |
| PGS*Grades of schooling |  |  |  | 0.006 |  |  |  | -0.008 |
|  |  |  |  | (.006) |  |  |  | (.006) |
| N | 5335 | 5335 | 5335 | 5335 | 5521 | 5521 | 5521 | 5521 |

*Notes:* The CES-D score is normalized to have a mean of 0 and standard deviation of 1. Grades of schooling is centered on 12 grades. All regressions include year of birth indicators, gender, and the first 10 principal components. Regressions in columns 3, 4, 7 and 8 additionally control for birth order, adolescent IQ, mother's schooling, an indicator for missing mother's schooling, and a PGS for educational attainment. Robust standard errors in parentheses. \*\*\*significant at 1% \*\*significant at 5% \*significant at 10%

Results for depression are given in table 3. There are 3 important observations. First, a 1 standard deviation increase in the PGS is associated with a 2.9 (2.3) percentage point increase in the probability of being depressed in Add Health (the WLS) in column 1 (5). The percentage effect relative to the sample mean is much smaller in Add Health (8%) than in the WLS (28%), but the percentage effect relative to the sample standard deviation is similar in both datasets (8% in Add Health and 7% in the WLS). Second, the schooling-depression gradient is 9 times larger in Add Health than in the WLS. Conditional on genetics, an extra grade of schooling is associated with a 2.6 (0.3) percentage point decrease in the likelihood of being depressed in Add Health (the WLS) in column 3 (7). Third, there is some suggestive evidence of schooling attenuating the genetic predisposition of depression at older ages but not in young adulthood. The GxE association is negative and statistically significant at the 5% level in the WLS, whereas it is positive and statistically insignificant in Add Health.

**Table 3:** OLS Associations of the PGS, Grades of Schooling, and GxE Interactions with Depression in Add Health and the WLS.

|  | Add Health |  |  |  | WLS |  |  |  |
| --- | --- | --- | --- | --- | --- | --- | --- | --- |
|  | (1) | (2) | (3) | (4) | (5) | (6) | (7) | (8) |
| **Outcome Mean** | 0.15 | 0.15 | 0.15 | 0.15 | 0.08 | 0.08 | 0.08 | 0.08 |
| **Genetics** |  |  |  |  |  |  |  |  |
| PGS | 0.029*** | 0.022*** | 0.023*** | 0.019** | 0.023*** | 0.022*** | 0.024*** |  |
| 0.032*** | (.005) | (.005) | (.005) | (.009) | (.004) | (.004) | (.128) | (.005) |
| **Schooling** |  |  |  |  |  |  |  |  |
| Grades of schooling |  | -0.029*** | -0.026*** | -0.026*** |  | -0.007*** | -0.003* | -0.003* |
|  |  | (.002) | (.003) | (.003) |  | (.002) | (.002) | (.002) |
| **GxE** |  |  |  |  |  |  |  |  |
| PGS*Grades of schooling |  |  |  | 0.002 |  |  |  | -0.005** |
|  |  |  |  | (.002) |  |  |  | (.002) |
| **N** | 5335 | 5335 | 5335 | 5335 | 5521 | 5521 | 5521 | 5521 |

*Notes:* All regressions include year of birth indicators, gender, and the first 10 principal components. Grades of schooling is centered on 12 grades. Regressions in columns 3, 4, 7 and 8 additionally control for birth order, adolescent IQ, mother's schooling, an indicator for missing mother's schooling, and a PGS for educational attainment. Robust standard errors in parentheses. \*\*\*significant at 1% \*\*significant at 5% \*significant at 10%

Table 4 presents results using a nonlinear specification for the PGS. In particular, we use dummy variables indicating whether the individual is in the 1^st^, 2^nd^, 3^rd^ (reference category), 4^th^, or 5^th^ quintile of the PGS distribution. The nonlinear PGS specifications show that individuals with a low (high) genetic predisposition have fewer (more) depressive symptoms relative to individuals in the middle of the PGS distribution in both datasets. For example, in Add Health the estimates in column 1 indicate that individuals in the 1^st^ quintile of the PGS distribution have a CES-D score that is 0.162 standard deviations lower than individuals in the 3^rd^ quintile. In contrast, individuals in the 5^th^ quintile have a CES-D score that is 0.122 standard deviations higher than individuals in the 3^rd^ quintile. The estimates in column 3 for the WLS show that the average difference in the CES-D score between individuals in the 1^st^ and 3^rd^ (5^th^ and 3^rd^) quintiles is −0.069 (0.177) standard deviations. However, the nonlinear specification gives different results for depression. In Add Health (column 5), there is a statistically significant difference (3.7 percentage points) in the probability of depression between individuals in the 1^st^ and 3^rd^ quintiles, but there is no statistically significant difference between individuals in the 5^th^ and 3^rd^ quintiles. For the WLS (column 7), there is a statistically significant difference in the likelihood of depression (6.2 percentage points) between individuals in the 5^th^ and 3^rd^ quintile, but no statistically significant difference between individuals in the 1^st^ and 3^rd^ quintiles. There is no evidence though that the association of grades of schooling with the CES-D score and depression significantly differs for individuals in the 1^st^ and 5^th^ quintile relative to individuals in the 3^rd^ quintile.

**Table 4:** OLS Associations of the PGS, Grades of Schooling, and GxE Interactions with Mental Health in Add Health and the WLS.

|  | CES-D Score |  |  |  | Depression |  |  |  |
| --- | --- | --- | --- | --- | --- | --- | --- | --- |
|  | Add Health<br>(1) | Add Health<br>(2) | WLS<br>(3) | WLS<br>(4) | Add Health<br>(5) | Add Health<br>(6) | WLS<br>(7) | WLS<br>(8) |
| **Genetics** |  |  |  |  |  |  |  |  |
| PGS 1<sup>st</sup> quintile | -0.162***<br>(.040) | -0.131*<br>(.073) | -0.069*<br>(.039) | -0.090*<br>(.053) | -0.037**<br>(.015) | -0.021<br>(.028) | -0.011<br>(.010) | -0.017<br>(.014) |
| PGS 2<sup>nd</sup> quintile | -0.078*<br>(.040) | 0.042<br>(.071) | -0.013<br>(.041) | -0.037<br>(.052) | -0.019<br>(.015) | 0.019<br>(.028) | 0.010<br>(.011) | -0.001<br>(.014) |
| PGS 4<sup>th</sup> quintile | 0.026<br>(.044) | 0.074<br>(.074) | 0.104**<br>(.042) | 0.110*<br>(.055) | 0.018<br>(.017) | 0.042<br>(.028) | 0.024**<br>(.012) | 0.029*<br>(.016) |
| PGS 5<sup>th</sup> quintile | 0.122**<br>(.044) | 0.144**<br>(.070) | 0.177***<br>(.044) | 0.176***<br>(.056) | 0.025<br>(.016) | 0.034<br>(.027) | 0.062***<br>(.013) | 0.074***<br>(.016) |
| **Schooling** |  |  |  |  |  |  |  |  |
| Grades of schooling | -0.085***<br>(.007) | -0.068***<br>(.015) | -0.007<br>(.006) | -0.011<br>(.012) | -0.026***<br>(.003) | -0.019***<br>(.005) | -0.003*<br>(.002) | -0.003<br>(.003) |
| **GxE** |  |  |  |  |  |  |  |  |
| PGS 1<sup>st</sup> quintile* |  | -0.012<br>(.020) |  | 0.011<br>(.016) |  | -0.006<br>(.007) |  | 0.003<br>(.004) |
| Grades of schooling |  |  |  |  |  |  |  |  |
| PGS 2<sup>nd</sup> quintile* |  | -0.045**<br>(.019) |  | 0.014<br>(.017) |  | -0.014*<br>(.007) |  | 0.006<br>(.005) |
| Grades of schooling |  |  |  |  |  |  |  |  |
| PGS 4<sup>th</sup> quintile* |  | -0.018<br>(.022) |  | -0.003<br>(.017) |  | -0.009<br>(.008) |  | -0.003<br>(.005) |
| Grades of schooling |  |  |  |  |  |  |  |  |
| PGS 5<sup>th</sup> quintile* |  | -0.007<br>(.020) |  | 0.000<br>(.018) |  | -0.003<br>(.007) |  | -0.008<br>(.005) |
| Grades of schooling |  |  |  |  |  |  |  |  |
| N | 5335 | 5335 | 5521 | 5521 | 5521 | 5521 | 5521 | 5521 |

*Notes:* The CES-D score is normalized to have a mean of 0 and standard deviation of 1. The indicator for the 3<sup>rd</sup> quintile of the PGS distribution is the reference category. Grades of schooling is centered on 12 grades. All regressions include year of birth indicators, gender, the first 10 principal components, birth order, adolescent IQ, mother's schooling, an indicator for missing mother's schooling, and a PGS for educational attainment. Robust standard errors in parentheses. \*\*\*significant at 1% \*\*significant at 5% \*significant at 10%

Table 5 presents results using an indicator variable for being a college graduate rather than grades of schooling. Using grades of schooling constrains the marginal impact of schooling attainment (as well as the one for the GxE interaction) to be the same over all schooling levels, but there may be larger associations at the upper end of the schooling distribution. With a college indicator, the GxE associations are still statistically insignificant in Add Health. However, in the WLS they are negative and statistically significant for both depressive symptoms and depression, indicating that the association of the PGS with mental health is smaller for college graduates than non-college graduates. The estimates in column 6 indicate that a 1 standard deviation increase in the PGS is associated with a 0.048 standard deviation increase in the CES-D score for college graduates, and a 0.111 standard deviation increase for non-college graduates. In column 8, a 1 standard deviation increase in the PGS is associated with a 3.2 percentage point increase in the probability of being depressed for non-college graduates, but only a 0.005 percentage point increase for college graduates. Appendix table A1 shows results from a regression the nonlinear PGS specification and the college graduate indicator. The GxE associations are statistically insignificant for both Add Health and the WLS. The one exception is a marginally statistically significant negative interaction between the college indicator and being in the 5^th^ quintile of the PGS distribution for depression in the WLS.

**Table 5:** OLS Associations of the PGS, College Graduation, and GxE Interactions with Mental Health in Add Health and the WLS.

|  | Add Health |  |  |  | WLS |  |  |  |
| --- | --- | --- | --- | --- | --- | --- | --- | --- |
|  | CES-D<br>(1) | CES-D<br>(2) | DEP<br>(3) | DEP<br>(4) | CES-D<br>(5) | CES-D<br>(6) | DEP<br>(7) | DEP<br>(8) |
| **Outcome Mean** | 0.00 | 0.00 | 0.15 | 0.15 | 0.00 | 0.00 | 0.08 | 0.08 |
| **Genetics** |  |  |  |  |  |  |  |  |
| PGS | 0.099***<br>(.013) | 0.102***<br>(.017) | 0.024***<br>(.005) | 0.025***<br>(.006) | 0.093***<br>(.014) | 0.111***<br>(.017) | 0.024***<br>(.004) | 0.032***<br>(.005) |
| **Schooling** |  |  |  |  |  |  |  |  |
| College graduate | -0.299***<br>(.030) | -0.301***<br>(.030) | -0.087***<br>(.011) | -0.087***<br>(.011) | -0.056*<br>(.031) | -0.059*<br>(.031) | -0.022***<br>(.008) | -0.024***<br>(.008) |
| **GxE** |  |  |  |  |  |  |  |  |
| PGS*College |  | -0.011<br>(.027) |  | -0.003<br>(.009) |  | -0.063**<br>(.028) |  | -0.027***<br>(.008) |
| N | 5335 | 5335 | 5335 | 5335 | 5521 | 5521 | 5521 | 5521 |

*Notes:* The CES-D score is normalized to have a mean of 0 and standard deviation of 1. All regressions include year of birth indicators, gender, the first 10 principal components, birth order, adolescent IQ, mother's schooling, an indicator for missing mother's schooling, and a PGS for educational attainment. Robust standard errors in parentheses. \*\*\*significant at 1% \*\*significant at 5% \*significant at 10%

Tables 6 and 7 show results from quantile regressions to investigate whether there are any heterogeneous associations along the conditional CES-D score distribution. We measure schooling attainment with grades of schooling in table 6, and with a college indicator in table 7. In both datasets, the association of the PGS is much larger in upper parts of the CES-D score distribution. For example, conditional on schooling attainment, a 1 standard deviation increase in the PGS is associated with a 0.106 standard deviation increase in the CES-D score in the 75^th^ conditional percentile (column 5), and a 0.211 standard deviation increase at the 90^th^ conditional percentile (column 7) in both datasets. In Add Health, the association of grades of schooling is also larger in upper parts of the conditional CES-D score distribution. In comparison, in the WLS there is no consistent increase in the (absolute) association of grades of schooling. However, in the WLS there is some suggestive evidence of schooling attainment moderating genetic predisposition. Specifically, there are negative and statistically significant GxE associations in the 75^th^ and 90^th^ percentile of the conditional CES-D score distribution. The GxE associations are also fairly large when using a college indicator to measure schooling attainment (table 7). In particular column 8 (6) in table 7 indicates that the marginal association of the PGS with the CES-D score is 0.222 (0.100) standard deviations smaller for college graduates at the 90^th^ (75^th^) percentile than for non-college graduates.^14^

**Table 6:** Quantile Regressions Associations of the PGS, Grades of Schooling and GxE interactions with the CES-D score in Add Health and the WLS.

|  | 25th<br>(1) | 25th<br>(2) | 50th<br>(3) | 50th<br>(4) | 75th<br>(5) | 75th<br>(6) | 90<sup>th</sup><br>(7) | 90<sup>th</sup><br>(8) |
| --- | --- | --- | --- | --- | --- | --- | --- | --- |
| **Panel A: Add Health** |  |  |  |  |  |  |  |  |
| **Genetics** |  |  |  |  |  |  |  |  |
| PGS | 0.060*** | 0.062*** | 0.092*** | 0.102*** | 0.106*** | 0.085*** | 0.211*** |  |
| 0.161*** | (.012) | (.018) | (.015) | (.027) | (.020) | (.028) | (.029) | (.052) |
| **Schooling** |  |  |  |  |  |  |  |  |
| Grades of schooling | -0.048*** | -0.048*** | -0.079*** | -0.081*** | -0.122*** | -0.121*** | -0.158*** | - |
| 0.151*** | (.006) | (.006) | (.008) | (.008) | (.011) | (.011) | (.018) | (.018) |
| **GxE** |  |  |  |  |  |  |  |  |
| PGS*Grades of schooling |  | -0.001<br>(.005) |  | -0.003<br>(.007) |  | 0.007<br>(.007) |  | 0.019<br>(.013) |
| N | 5335 | 5335 | 5335 | 5335 | 5335 | 5335 | 5335 | 5335 |
| **Panel B: WLS** |  |  |  |  |  |  |  |  |
| **Genetics** |  |  |  |  |  |  |  |  |
| PGS | 0.005*** | 0.041*** | 0.075*** | 0.078*** | 0.106*** | 0.148*** | 0.222*** |  |
| 0.248*** | (.010) | (.013) | (.013) | (.017) | (.015) | (.019) | (.029) | (.036) |
| **Schooling** |  |  |  |  |  |  |  |  |
| Grades of schooling | -0.001<br>(.004) | -0.000<br>(.005) | 0.002<br>(.008) | 0.003<br>(.006) | -0.000<br>(.007) | -0.001***<br>(.007) | -0.020***<br>(.014) | -0.028*<br>(.014) |
| **GxE** |  |  |  |  |  |  |  |  |
| PGS*Grades of schooling |  | -0.003<br>(.004) |  | -0.002<br>(.005) |  | -0.019***<br>(.007) |  | -0.022*<br>(.013) |
| N | 5521 | 5521 | 5521 | 5521 | 5521 | 5521 | 5521 | 5521 |

Notes: The CES-D score is normalized to have a mean of 0 and standard deviation of 1. All regressions include year of birth indicators, gender, the first 10 principal components, birth order, adolescent IQ, mother's schooling, an indicator for missing mother's schooling, and a PGS for educational attainment. Robust standard errors in parentheses. \*\*\*significant at 1% \*\*significant at 5% \*significant at 10%

**Table 7:**
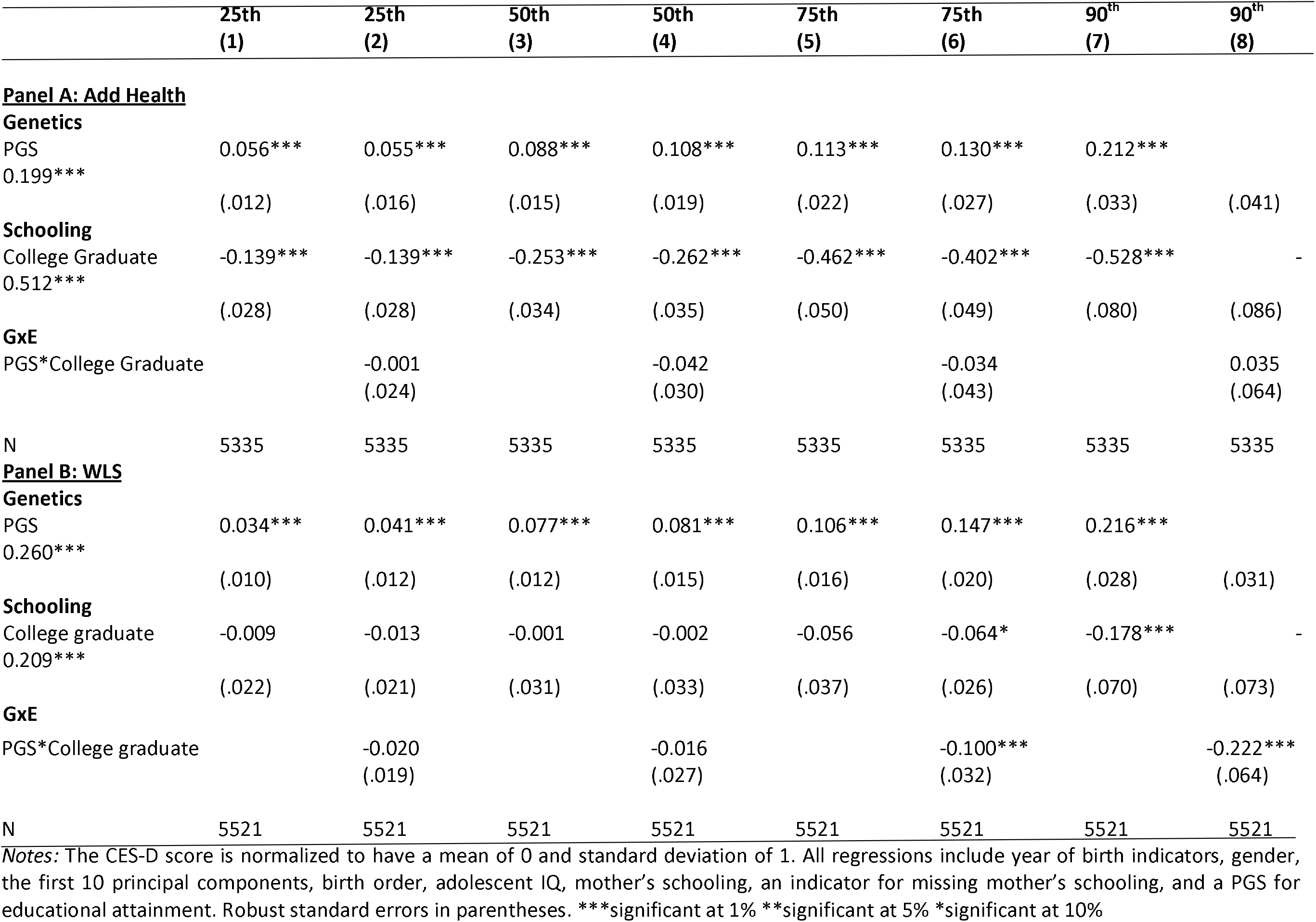
Quantile Regressions Associations of the PGS, College Graduation and GxE interactions with the CES-D score in Add Health and the WLS.

Overall, the OLS results show that there are large associations of the PGS with mental health, and large differences in mental health by schooling attainment in both datasets. There is some suggestive evidence of GxE interactions in the WLS, but no consistent evidence in Add Health. However, since there may be unobserved confounders left uncontrolled, the OLS estimates may be biased. We now turn to the sibling fixed-effects analyses, which control for unobserved family-level factors and about 50% of unobserved genetic factors that might bias the OLS estimates.

### 4.3 Sibling Fixed-Effects Results

OLS and sibling fixed-effects results for the CES-D score are shown in table 8. Panel A presents results using grades of schooling and panel B shows results using a college graduate indicator. The OLS and sibling fixed-effects estimates in columns 1 and 3 show that conditional on schooling attainment the PGS is not statistically significantly associated with depressive symptoms in Add Health. In comparison the OLS estimate for the WLS in column 5 shows that a 1 standard deviation increase in the PGS is associated with a 0.096 standard deviation increase in the CES-D score conditional on schooling attainment. The sibling fixed-effects estimate in column 7 is slightly larger at 0.107. Both the OLS and sibling fixed-effects estimates are similar to the corresponding OLS estimate of 0.093 for the full sample in table 2 column 7. In terms of schooling attainment the OLS results show that higher schooling attainment is associated with fewer depressive symptoms in Add Health (column 2) but not in the WLS (column 5). The sibling fixed-effects results in column 3 and 7 show that grades of schooling are not significantly associated with depressive symptoms in Add Health and the WLS. However, being a college graduate is still significantly associated with a lower CES-D score, even when controlling for unobserved family and shared genetic factors. Moreover, the magnitude of the sibling fixed-effects estimate (0.325) for the college graduate indicator is almost the same as the corresponding OLS estimate (0.334) in Add Health, and it is also similar to that for the corresponding estimate for the full sample in table 4 column 1 (0.299). In the WLS, the sibling fixed-effects estimate (0.125) is twice the size of the OLS estimate (0.050). In both datasets there is no strong evidence of any GxE associations.

**Table 8:** OLS and Sibling Fixed-Effects Associations of the PGS, Schooling Attainment, and GxE interactions with the CES-D Score in Add Health and the WLS.

|  | Add Health |  |  |  | WLS |  |  |  |
| --- | --- | --- | --- | --- | --- | --- | --- | --- |
|  | OLS<br>(1) | OLS<br>(2) | FE<br>(3) | FE<br>(4) | OLS<br>(5) | OLS<br>(6) | FE<br>(7) | FE<br>(8) |
| **Panel A: Grades of Schooling** |  |  |  |  |  |  |  |  |
| **Genetics** |  |  |  |  |  |  |  |  |
| PGS | 0.042<br>(.033) | 0.051<br>(.046) | 0.011<br>(.058) | 0.030<br>(.089) | 0.096***<br>(.022) | 0.113***<br>(.128) | 0.107***<br>(.040) | 0.116**<br>(.046) |
| **Schooling** |  |  |  |  |  |  |  |  |
| Grades of schooling | -0.056***<br>(.017) | -0.056***<br>(.017) | -0.013<br>(.029) | -0.012<br>(.029) | -0.003<br>(.010) | -0.003<br>(.010) | -0.007<br>(.015) | -0.007<br>(.010) |
| **GxE** |  |  |  |  |  |  |  |  |
| PGS*Grades of schooling | --- | -0.004<br>(.013) | --- | -0.008<br>(.025) | --- | -0.009<br>(.009) | --- | -0.005<br>(.012) |
| **Panel B: College Graduate** |  |  |  |  |  |  |  |  |
| **Genetics** |  |  |  |  |  |  |  |  |
| PGS | 0.035<br>(.032) | 0.049<br>(.041) | 0.008<br>(.058) | 0.035<br>(.072) | 0.096***<br>(.022) | 0.121***<br>(.027) | 0.104***<br>(.040) | 0.127***<br>(.045) |
| **Schooling** |  |  |  |  |  |  |  |  |
| College graduate | -0.334***<br>(.076) | -0.338***<br>(.076) | -0.325***<br>(.109) | -0.328***<br>(.110) | -0.050<br>(.046) | -0.054<br>(.045) | -0.125*<br>(.069) | -0.125*<br>(.068) |
| **GxE** |  |  |  |  |  |  |  |  |
| PGS*College graduate | --- | -0.044<br>(.058) | --- | -0.079<br>(.090) | --- | -0.077*<br>(.044) | --- | -0.073<br>(.062) |
| N | 914 | 914 | 914 | 914 | 2228 | 2228 | 2228 | 2228 |

*Notes:* The CES-D score is normalized to have a mean of 0 and standard deviation of 1. Grades of schooling is centered on 12 grades. All regressions include year of birth indicators, gender, the first 10 principal components, birth order, adolescent IQ, mother's schooling, an indicator for missing mother's schooling, and a PGS for educational attainment. Robust standard errors clustered by sibling pairs in parentheses. \*\*\*significant at 1% \*\*significant at 5% \*significant at 10%

Results for depression are presented in table 9. In the WLS, both the OLS and sibling fixed-effects estimates show that the PGS is significantly associated with a higher probability of depression. The OLS estimates show that college graduates are less likely to be depressed than non-college graduates, but this association becomes statistically insignificant in the sibling fixed-effects models. The GxE associations are negative but statistically insignificant in both the OLS and sibling fixed-effects models. Note that, in general, the OLS estimates for siblings in the WLS in columns 5 and 6 of table 9 are similar in magnitude to the corresponding ones for the full sample in columns 7 and 8 of tables 3 and 4. For Add Health the OLS estimates only show a significant association of schooling attainment, with all the sibling fixed-effects estimates being statistically insignificant.

**Table 9:**
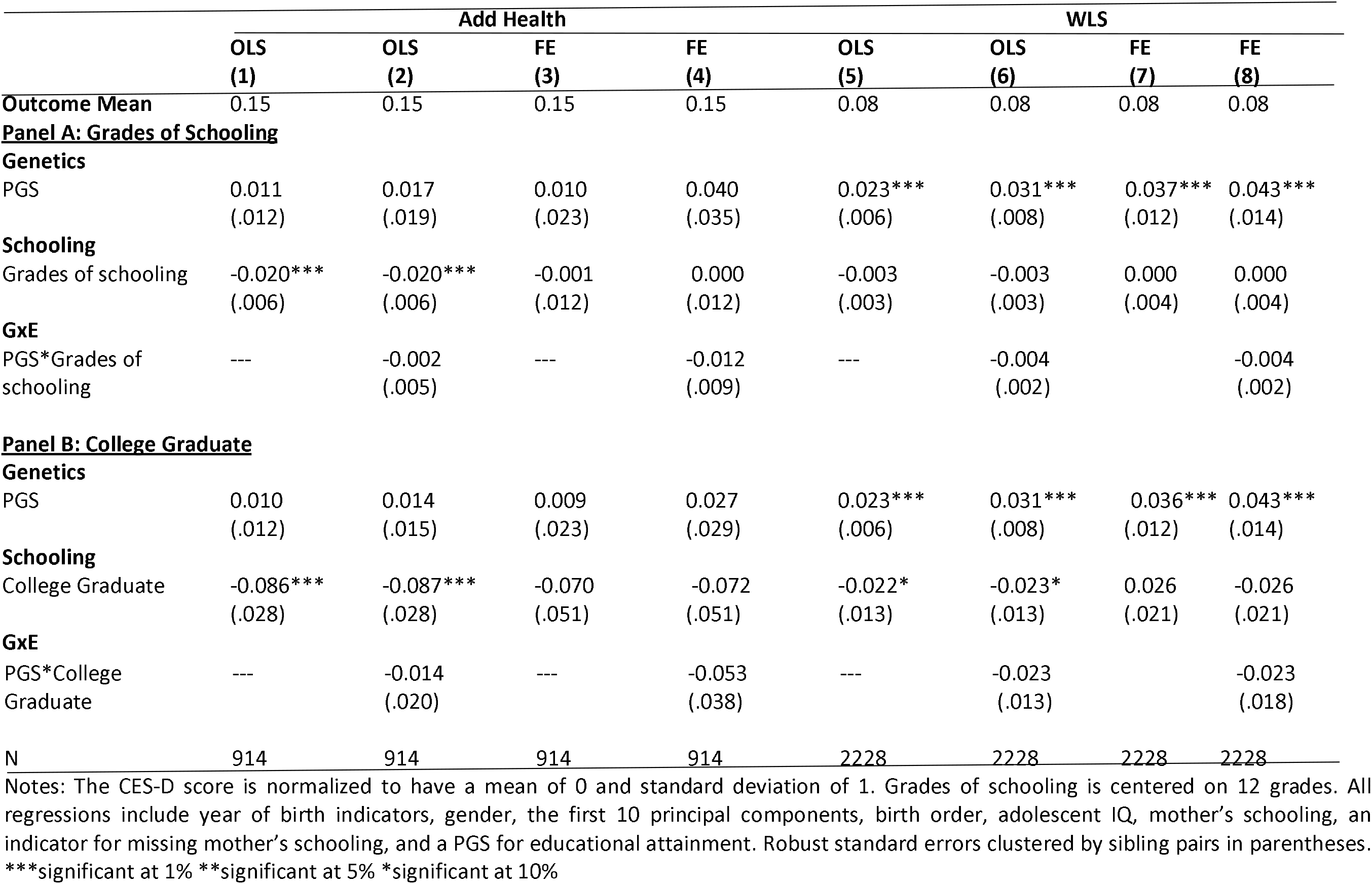
OLS and Sibling Fixed-Effects Associations of the PGS, Schooling attainment, and GxE interactions with the Depression in Add Health and the WLS.

It is difficult to draw firm conclusions about GxE interactions from the sibling fixed-effects estimates in Add Health, because the PGS is not associated with mental health in OLS models. One possible reason for this could be that the Add Health sibling pairs are a selected sample. However, the descriptive statistics suggested that the sibling pairs are similar in observed characteristics to individuals in the main estimation sample. We checked whether there are any statistically significant differences in schooling attainment, mental health, and background characteristics between the Add Health sibling pair sample and the Add Health estimation sample excluding sibling. Appendix table A2 shows that there are no statistically significant differences in schooling attainment or mental health.^15^ There is however a statistically significant difference in adolescent IQ, with siblings have a slight lower IQ score. Nevertheless, the OLS estimates show statistically significant associations of schooling attainment, which are similar in magnitude to those for the main estimation sample. For example, in table 8 column 1 an extra grade of schooling is associated with a 0.056 standard deviation decrease in the CES-D score conditional on the PGS. The corresponding association in table 2 column 3 for the main estimation sample is 0.085. Another possible reason could be low statistical power given the sample size is relatively small. In the WLS, the PGS is associated with mental health but the GxE associations are statistically insignificant. This could also be due to low power. We note that the point estimates of the GxE associations in the sibling fixed-effects models are similar in magnitude to the GxE associations in the OLS analyses for the main estimation sample. For example, in the OLS analyses the GxE association (standard error) for depressive symptoms was −0.063 (0.028) in table 4 column 6. The sibling fixed-effects estimate in table 8 column 8 is similar in magnitude (−0.073) but the standard error is almost double (0.062). Similarly, the sibling fixed-effects GxE estimate for depression (−0.023; table 9 column 8) is similar to the OLS estimate for the full sample (−0.027; table 4 column 8), but the higher standard error renders it insignificant. Finally, although we cannot draw strong conclusions about GxE interactions, there is an interesting pattern of results for schooling attainment. In both datasets, we find that a college education is associated with fewer depressive symptoms even when controlling for shared family characteristics, but there is no association for grades of schooling.

## 5. Conclusions

The diathesis-stress and the differential susceptibility theories for genetic influences on health suggest that higher schooling attainment may attenuate the genetic predisposition to poor health. A few studies have investigated this hypothesis for BMI and obesity (Amin et al. 2017; Liu et al; 2015; Komulainen et al. 2018). OLS results from these studies show statistically insignificant GxE associations. We contribute by examining this hypothesis for mental health in two distinct US datasets (Add Health and the WLS) at different adult ages. To our knowledge, this is the first study to provide estimates of the associations of genetics, schooling attainment, and GxE interactions with mental health.

Our OLS results indicate that the association of the PGS with mental health is similar in Add Health and the WLS, but the association of schooling attainment is much larger in Add Health than in the WLS. That the schooling-mental health gradient is smaller in the WLS is not surprising because the majority of respondents in the WLS are high school graduates. There is some suggestive evidence that the association of the PGS with mental health is lower for more-schooled older individuals in the WLS, but there is no evidence of any significant GxE interactions in Add Health. Our results from quantile regressions indicate that the GxE interactions are larger and statistically significant in the upper part of the conditional CES-D score distribution. We assess the robustness of the OLS results to potential omitted variable bias by using the siblings samples in both datasets to estimate sibling fixed-effect regressions. It is difficult to draw firm conclusions from the sibling fixed-effects results, in part due to low statistical power. However, the sibling fixed-effects estimates show that college education is associated with fewer depressive symptoms in both datasets. Thus, the evidence differs between the two data sets, with fairly robust results concerning significant associations between PGS and schooling attainment and mental health, as well as suggestive evidence for protective effects of more schooling, for the WLS -- but not robust significant results for PGS or for GxE interactions for Add Health. Given the different ages of the two groups, this raises the question of whether the associations with PGS and schooling-PGS interactions are revealed only in mature adulthood. This suggests the value of future research to investigate the age-dependence of such associations over a broader age range.

## Footnotes

1 Border et al. (2019) use the UK Biobank to look at candidate GxE associations across several measures of depression (lifetime diagnosis, current severity, episode recurrence) and environmental factors (sexual or physical abuse during childhood, socioeconomic adversity). Their study is well-powered with sample sizes ranging from 62,138 to 443,264. Despite the large sample sizes, they find no clear evidence for any candidate GxE associations.

2 Mullins et al. (2016) explain this result by suggesting that childhood trauma is such a strong risk factor for MDD that it overrides some of the genetic risk of MDD.

3 There is also some evidence that there is a causal relationship between schooling and mental health. See for example Crespo et al. (2014) and Mazzonna (2014). Both studies find evidence for a causal effect of schooling by exploiting variation in schooling arising from changes in compulsory education laws, which can be argued to be exogenous. However, other studies using the same methodology have found no protective effect of schooling on mental health (Avendano et al. 2017; Courtin et al. 2018; Dursuin and Cesur 2016)

4 We limit the analyses to European-ancestry individuals to control for biases due to population stratification. This is explained in more detail in section 3.

5 The results from these studies cannot be given a causal interpretation because of omitted variable bias. A study that address omitted variable bias is Barcellos et al. (2018). They use the raising of the minimum school leaving age in the UK to form instruments for schooling attainment. Their instrumental variable results show robust and statistically significant negative GxE associations between schooling attainment and the PGS for BMI.

6 See Figure 1 in Galama and van Kippersluis (2013), which shows that socioeconomic health gradients exist by age 20. The gradient increases until about age 60, after which it narrows.

7 We do not use wave 5 as the data has not been released.

8 Detailed information about the construction of the PGS is given by Add Health documentation at: https://www.cpc.unc.edu/projects/addhealth/documentation/guides/SSGACPGS_UsersGuide.pdf

9 The key assumption is that all SNPs share the same variance-covariance matrix of effect sizes across traits. This assumption is violated, for example, if some SNPs influence only a subset of the traits. Turley et al. (2018) analytically show that MTAG is still a consistent estimator, and that MTAG PGSs outperform PGSs based on standard GWAS summary statistics, even if this assumption is violated.

10 Of the 9,897 individuals, 4,187 are of non-European ancestry and were dropped by the SSGAC. The SSGAC additionally excluded individuals who did not satisfy the following criteria: (i) genotype missingness rate is less than 0.05 in all chromosomes, (ii) there is no mismatch between surveyed sex and genetic sex, (iii) the individual is not an outlier in terms of heterozygosity/homozygosity, and (iv) the individual is not an ancestral outlier. SNPs were also dropped with a call rate less than 0.98, Hardy-Weinberg exact test *P*-value less than 10^-4^, or minor allele frequency > 0.01.

11 At wave 1, there are 20,745 individuals, and mother’s schooling is missing for 11% (2332 individuals).

12 Add Health implemented an abridged (87 items) version of the Peabody Picture Vocabulary Test (PPVT) - Revised. In the test, the interviewer reads a word aloud and the respondent selects the illustration that best fits its meaning. For example, the word “furry” has illustrations of a parrot, dolphin, frog, and cat from which to choose. The test proceeds through a series of rounds with increasing difficulty. Test scores are standardized by age with mean 100.

13 Compared to Add Health, we lose considerably more observations in the WLS. This is because the CES-D score is only available for 6,876 respondents that also have data on the PGS.

14 We tested whether the GxE coefficient estimates from the quantile regressions are statistically different from each other. In the WLS, we rejected the null hypothesis that the GxE coefficients (when using the college indicator) are equal across quantiles. We could not reject the null hypothesis of the equality of GxE coefficients across quantiles for the WLS when using grades of schooling, and for Add Health.

15 We also performed the Kolmogornov-Smirnov test for equality of distributions. We could not reject the null hypothesis that the CES-D score distribution, grades of schooling distribution, and PGS distribution between the Add Health sibling sample and Add Health main estimation sample (excluding siblings) are equal.

---

**Appendix Table A1:** OLS Associations of the PGS, College Graduation, and GxE Interactions with Mental Health in Add Health and the WLS.

|  | CES-D Score |  |  |  | Depression |  |  |  |
| --- | --- | --- | --- | --- | --- | --- | --- | --- |
|  | Add Health<br>(1) | Add Health<br>(2) | WLS<br>(3) | WLS<br>(4) | Add Health<br>(5) | Add Health<br>(6) | WLS<br>(7) | WLS<br>(8) |
| **Genetics** |  |  |  |  |  |  |  |  |
| PGS 1<sup>st</sup> quintile | -0.164***<br>(.040) | -0.175***<br>(.055) | -0.067*<br>(.039) | -0.092*<br>(.050) | -0.038**<br>(.015) | -0.042**<br>(.020) | -0.011<br>(.011) | -0.018<br>(.013) |
| PGS 2<sup>nd</sup> quintile | -0.078*<br>(.041) | -0.053<br>(.055) | -0.012<br>(.041) | -0.042<br>(.050) | -0.019<br>(.015) | -0.011<br>(.020) | -0.010<br>(.011) | 0.003<br>(.014) |
| PGS 4<sup>th</sup> quintile | 0.025<br>(.044) | 0.033<br>(.056) | 0.105**<br>(.042) | 0.120**<br>(.052) | 0.018<br>(.016) | 0.023<br>(.021) | 0.024**<br>(.012) | 0.029*<br>(.015) |
| PGS 5<sup>th</sup> quintile | 0.123***<br>(.044) | 0.134***<br>(.056) | 0.177***<br>(.044) | 0.181***<br>(.054) | 0.026<br>(.016) | 0.030<br>(.021) | 0.062***<br>(.013) | 0.073***<br>(.016) |
| **Schooling** |  |  |  |  |  |  |  |  |
| College graduate | -0.300***<br>(.030) | -0.280***<br>(.078) | -0.059*<br>(.031) | -0.083<br>(.058) | -0.087***<br>(.011) | -0.079***<br>(.017) | -0.023***<br>(.008) | -0.020<br>(.016) |
| **GxE** |  |  |  |  |  |  |  |  |
| PGS 1st quintile* |  | 0.024 |  | 0.075 |  | -0.010 |  | 0.020 |
| College graduate |  | (.078) |  | (.079) |  | (.028) |  | (.021) |
| PGS 2<sup>nd</sup> quintile* |  | -0.070 |  | 0.100 |  | -0.022 |  | 0.022 |
| College graduate |  | (.078) |  | (.085) |  | (.029) |  | (.023) |
| PGS 4<sup>th</sup> quintile* |  | -0.023 |  | -0.048 |  | -0.015 |  | -0.018 |
| College graduate |  | (.088) |  | (.085) |  | (.032) |  | (.023) |
| PGS 5<sup>th</sup> quintile* |  | -0.036 |  | -0.017 |  | -0.014 |  | -0.043* |
| College graduate |  | (.088) |  | (.090) |  | (.032) |  | (.025) |
| N | 5335 | 5335 | 5521 | 5521 | 5335 | 5335 | 5521 | 5521 |

*Notes:* The CES-D score is normalized to have a mean of 0 and standard deviation of 1. . The indicator for the 3<sup>rd</sup> quintile of the PGS distribution is the reference category. All regressions include year of birth indicators, gender, the first 10 principal components, birth order, adolescent IQ, mother's schooling, an indicator for missing mother's schooling, and a PGS for educational attainment. Robust standard errors in parentheses.
\*\*\*significant at 1% \*\*significant at 5% \*significant at 10%

**Appendix Table A2:**
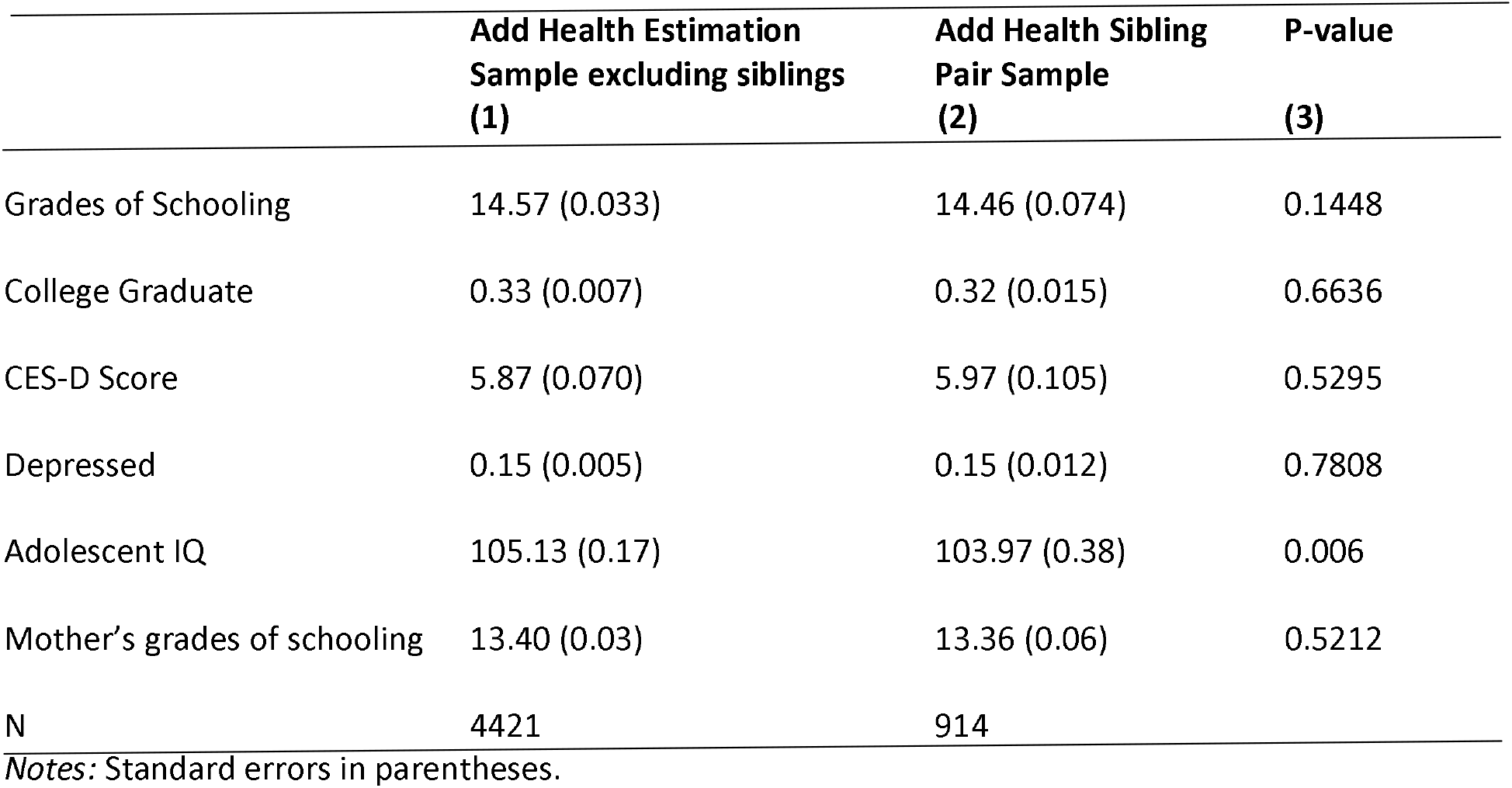
Comparison of Add Health Sibling Pair Sample and Add Health Main Estimation Sample Excluding Siblings.

